# Spike protein binding prediction with neutralizing antibodies of SARS-CoV-2

**DOI:** 10.1101/2020.02.22.951178

**Authors:** Tamina Park, Sang-Yeop Lee, Seil Kim, Mi Jeong Kim, Hong Gi Kim, Sangmi Jun, Seung Il Kim, Bum Tae Kim, Edmond Changkyun Park, Daeui Park

## Abstract

Coronavirus disease 2019 (COVID-19) is a new emerging human infectious disease caused by Severe Acute Respiratory Syndrome Coronavirus 2 (SARS-CoV-2, also previously known as 2019-nCoV), originated in Wuhan seafood and animal market, China. Since December 2019, more than 69,000 cases of COVID-19 have been confirmed in China and quickly spreads to other counties. Currently, researchers put their best efforts to identify effective drugs for COVID-19. The neutralizing antibody, which binds to viral capsid in a manner that inhibits cellular entry of virus and uncoating of the genome, is the specific defense against viral invaders. In this study, we investigate to identify neutralizing antibodies that can bind to SARS-CoV-2 Sipke (S) protein and interfere with the interaction between viral S protein and a host receptor by bioinformatic methods. The sequence analysis of S protein showed two major differences in the RBD region of the SARS-CoV-2 S protein compared to SARS-CoV and SARS-CoV related bat viruses (btSARS-CoV). The insertion regions were close to interacting residues with the human ACE2 receptor. Epitope analysis of neutralizing antibodies revealed that SARS-CoV neutralizing antibodies used conformational epitopes, whereas MERS-CoV neutralizing antibodies used a common linear epitope region, which contributes to form the β-sheet structure in MERS-CoV S protein and deleted in SARS-CoV-2 S protein. To identify effective neutralizing antibodies for SARS-CoV-2, the binding affinities of neutralizing antibodies with SARS-CoV-2 S protein were predicted and compared by antibody-antigen docking simulation. The result showed that CR3022 neutralizing antibody from human may have higher binding affinity with SARS-CoV-2 S protein than SARS-CoV S protein. We also found that F26G19 and D12 mouse antibodies could bind to SARS-CoV S protein with high affinity. Our findings provide crucial clues towards the development of antigen diagnosis, therapeutic antibody, and the vaccine against SARS-CoV-2.

## Introduction

*Coronaviridae* is a family of enveloped viruses which have a single strand, positive-stranded RNA genome and classified into four genera: ɑ, β, γ, and δ. Coronavirus (CoV) has been identified in human and animals including bats, camels, zpigs, cats, and mice. The viruses usually cause mild to moderate upper-respiratory tract illnesses in human [1]. Two of betacoronaviruses, severe acute respiratory syndrome (SARS) coronavirus (SARS-CoV) and Middle East respiratory syndrome (MERS) coronavirus (MERS-CoV), caused severe epidemics during the last two decades. SARS-CoV emerged in November 2002 in Guangdong province, China and affected 29 countries. The epidemic of SARS-CoV resulted in 8,096 human infections and 774 deaths (9.6% fatality rate) by July 2003 [2]. MERS-CoV was first reported in Saudi Arabia in September 2012 and spread to 28 countries. The epidemic of MERS-CoV resulted in 2,494 human infections and 858 deaths (34.4% fatality rate) by November 2019 [3].

Coronavirus disease 2019 (COVID-19) is newly emerging human infectious disease caused by Severe Acute Respiratory Syndrome Coronavirus 2 (SARS-CoV-2, also previous known as 2019-nCoV) originated in Wuhan seafood and animal market. In December 2019, a series of pneumonia cases of unknown cause have been reported in Wuhan, Hubei province, China [4]. Later, on January 7, a novel CoV was identified from the bronchoalveolar lavage fluid of a patient [5], and named SARS-CoV-2 by International Committee on Taxonomy of Viruses [6]. The early COVID-19 patients show symptoms of severe acute respiratory infection, such as fever, cough, sore throat, nasal congestion, headache, muscle pain or malaise, and severe patients develop to acute respiratory distress syndrome, sepsis or septic shock [7–9]. As of February 16, 2020, more than 69,200 cases of COVID-19 have been confirmed in China and quickly spreads to other counties [10].

The SARS-CoV-2 is a member of betacoronavirus and shows 79% and 50% sequence identity with SARS-CoV and MERS-CoV, respectively. Phylogenetic analysis revealed that SARS-CoV-2 is most similar (88% sequence identity) to SARS-like CoVs previously collected from bats in China [5, 11, 12]. Although the viral pathogenesis of SARS-CoV-2 is unknown, most recent reports suggest that SARS-CoV-2 may use angiotensin-converting enzyme II (ACE2) as a cellular entry receptor. ACE2 is a well known host cell receptor for SARS-CoV [12]. Shi and colleges showed that SARS-CoV-2 uses ACE2 as a cellular entry receptor but not other CoV receptors, aminopeptidase N (APN) and dipeptidyl peptidase 4 (DPP4) [13]. Ying and colleges showed the receptor-binding domain (RBD) of SARS-CoV-2 spike glycoprotein (S protein) interacts with ACE2 [14]. McLaellen and colleges showed that ACE2 binds to SARS-CoV-2 S protein with much higher affinity than to SARS-CoV S protein [15]. In addition, bioinformatic analysis proposed binding structure of RBD of S protein (S-RBD) and ACE2 [16]. Thus, it is of great interest to identify neutralizing antibodies that can interact with SARS-CoV-2 S-RBD and interfere with the binding between viral S protein and host receptor ACE2.

After the severe epidemic events of SARS and MERS, researchers have been made great efforts to discover neutralizing antibodies for CoVs [17, 18]. The neutralizing antibodies for CoVs mainly targeted to S-RBD. S protein of SARS-CoV-2 shows 76.2% and 34.1% amino acid sequence identity to those of SARS-CoV and MERS-CoV, respectively. Therefore, the neutralizing antibodies of SARS-CoV and MERS-CoV S proteins may have a possibility to interact with SARS-CoV-2 S protein and show similar viral neutralization effect. In the present study, we employed a antibody-antigen docking approach to predict the interaction between SARS-CoV-2 S-RBD and previously reported neutralization antibodies for SARS-CoV and MERS-CoV.

## Methods

### Phylogenetic analysis of SARS-CoV-2 S protein

To comparing of S gene containing S protein among SARS-CoV-2, SARS-CoV, and MERS-CoV strains, the nucleotide sequences of S gene were retrieved from GISAID [19] and ViPR [20]. The S genes of SARS-CoV-2 were retrieved from initially sequenced 62 genomes of SARS-CoV-2 strains. After removal of identical S gene sequences, 16 genes of S protein were used in the study. The sequence from SARS-CoV-2 Wuhan-Hu-1 (Genbank MN908947.3 or GISAID EPI ISL 402125) was used as a representative sequence of SARS-CoV-2 strains throughout this study. The closely related strains of SARS-CoV-2 were selected from preliminary and extensive phylogenetic analaysis of SARS-CoV related strains including btSARS-CoV, SARS-CoV and a MERS-CoV strain. More detail information of the sequences used in this study were listed in Supplementary Table S1. The sequence alignments and phylogenetic analysis were done using MEGA X [21]. The nucleotide sequence were codon aligned using ClustalW with default parameters and the phylogenetic tree was inferred using neighbor-joining [22], maximum-likelihood [23], and maximum-parsimony methods [24]. The distance matrix was calculated based on the Jukes-Cantor methods [25]. The bootstrap values of the phylogenetic tree were derived from 1,000 replicates [26].

### Conservation score and epitope mapping of SARS-CoV-2 S protein

The conservation score of amino acid positions on S protein in SARS-CoV-2 was calculated by ConSurf program [27]. The multiple sequence alignment of SARS-CoV-2 strain Wuhan-Hu-1and 12 related strains (Supplementary Table S1) was used as an input for ConSurf. In ConSurf, conservation scores and confidence intervals for the conservation scores were calculated using the empirical Bayesian method. The scores were normalized using the number of inputted sequences. Also, the highest score of ConSurf program means the most conserved position among sequences. We additionally checked the epitope positions on the SARS-CoV-2 S protein based on the known epitope information of 11 neutralizing antibodies developing for SARS-CoV and MERS-CoV. Each information of epitope positions was acquired from literatures (Table 1).

**Table 1.** Neutralizing antibodies and their epitopes analyzed in the study.

| Target | Name | Origin | RBD binding residues | PDB ID | Reference |
| --- | --- | --- | --- | --- | --- |
| SARS-CoV | m396 | Human | K365, K390, D392, R395, R426, D429, R441, E452, D454, F483, Y484, T485, T486, T487, G488, I489, Y491, Q492 | 2DD8 | [39] |
|  | 80R | Human | T433, N437, K439, P469, P470, A471, C474, W476, L478, D480, T485, Q492 | 2GHW | [40] |
|  | S230 | Human | L443, Y408, Y442, F460, Y475 | 6NB6 | [41] |
|  | F26G19 | Mouse | T486, T487, G488, I489, G490, Y491, Q492 | 3NGF | [42] |
|  | CR3022 | Human | n.a. | n.a. | n.a. |
| MERS-CoV | MERS-27 | Human | T392, N398, L495, K496, V527, S528, I529, V530, P531, S532, T533, W535, E536, D537, G538, D539, Y540, Y541, R542, K543, W553 | 5YY5 | [43, 44] |
|  | CDC2-C2 | Human | N501, K502, S504, F506, D510, R511, T512, E513, W535, E536, D537, G538, D539, Y540, Y541, R542, W553, V555, A556, S557 | 6C6Z | [43] |
|  | m336 | Human | Y499, N501, K502, S504, R505, F506, D510, R511, T512, E513, P515, W535, E536, D537, G538, D539, Y540, Y541, R542, W553, V555, S557, G558, S589 | 4XAK | [43, 45] |
|  | MCA1 | Human | K502, F506, S508, D510, R511, E513, S528, I529, P531, T533, W535, E536, G538, D539, Y540, Y541, R542, K543, Q544, W553, V555, S557 | 5GMQ | [43, 46] |
|  | 4C2 | Humanized | G391, T392, P394, Y397, N398, K400, L495, K496, Y523, P525, C526, V527, S528, I529, V530, P531, S532, T533, V534, W535, E536, D537, D539, Y540, Y541, R542, K543, Q544, L545, S546, E549, W553, T560 | 5DO2 | [43, 47] |
|  | D12 | Mouse | G391, T392, P394, Y397, N398, F399, K400, L495, K496, P525, V527, S528, I529, P531, S532, T533, W535, E536, D537, D539, Y540, Y541, R542, K543, Q544, L545, S546, E549, W553, T560 | 4ZPT | [43, 48] |

### Structure of SARS-CoV-2 S protein

S-RBD protein structure was used cryo-EM structure (Protein Data Bank ID: 6VSB) [15]. To predict the missing region of cryo-EM structure in SARS-CoV-2 S-RBD, we performed homology modeling based on known the three dimensional structure of SARS-CoV (PDB ID: 6NB7) using SWISS-MODEL (https://swissmodel.expasy.org/) [28]. Then, the best homology models were selected according to Qualitative Model Energy ANalysis (QMEAN) statistical parameter. The structures were visualized with UCSF’s Chimera (https://www.cgl.ucsf.edu/chimera/).

### Neutralizing antibody candidates

As neutralizing antibody candidates of SARS-CoV-2, the five antibodies against SARS-CoV and the six antibodies to prevent MERS-CoV were selected in the study (Table 1). The complex structure of RBD and ten neutralizing antibodies was retrieved from PDB. The complex structures were superimposed to the RBD structure of SARS-CoV-2 which were built by homology modeling. The procedures were performed that the RBD structures of SARS-CoV2 and SARS-CoV were aligned by pairwise sequence alignment. And then the structures were superimposed according to those pairwise alignments using MatchMaker program [29]. Finally, we suceesfully predicted the complex structures of neutralizaing antibody candidates and RBD of SARS-CoV-2. About the antibody such as CR3022 [30] that the structure was not revealed, we performed the antibody strcture modeling with Rosetta program [31].

### Antibody-antigen docking simulation

Docking simlutation between the RBD of SARS-CoV-2 and certain SARS-CoV and MERS-CoV antibodies were implemented with Rosetta antibody-antigen docking protocols [32]. Rosetta SnugDock program can refine homology models with the flexible and uncertain region, because the program simulates most of conformation space available to antibody paratopes [33]. With the complex structures of RBD and antibody candidates, all-atom relax protocol, docking prepack protocol, and antibody-antigen docking simulation were carried out to calculate the free energy of low-energy binding conformations. The distiribution of docking scores displayed as funnel plots using interface RMD (interface RMS) versus the binding score (dG binding) between antibody and antigen (Fig. 4). The binding score was used Rosetta’s docking interface score (based on the Talaris2013 force field) to rank the complexes. Rosetta interface score is defined as Isc = *E*bound − *E*unbound, where *E*bound is the score of the bound complex and *E*unbound is the sum of the scores of the individual protein partners in isolation. In addition, 1000 independent docking runs were performed to generate the antibody-antigen models. To predict possible neutralizing antibody candidates of SARS-CoV-2, the docking results were compared between interface binding scores of SARS-CoV-2 S-RBD (homology modeling) and interface binding scores of SARS-CoV or MERS-CoV S-RBD (crystal structure) with 11 antibodies for SARS-CoV and MERS-CoV have been developed. The statistical significance was tested using student’s *t-test*.

## Results and Discussion

### Phylogenetic analysis and amino acid variation of S protein

The phylogenetic tree showed that the protein gene sequences were clearly clustered into three groups; SARS-CoV-2 related, SARS-CoV related and HKU3 related groups (Fig. 1A). SARS-CoV and SARS-CoV-2 groups formed a rigid monophyletic group with their own closest bat SARS-CoV related strains, respectively. The result suggested that these two human-pathogenic CoV strains were derived from common ancestral bat CoV. The sequence alignments showed that there were insertions and deletion during the divergence among the strains (Fig. 1B and 1C). Various deletions were observed in SARS-CoV related group. The NTD region (position 71-77, GTNGTKR) of S protein in the strain Wuhan-Hu-1 was mostly conserved in SARS-CoV-2 group but not in SARS-CoV related group. The NTD region was also conserved in other btSARS-CoV strains but the sequence similarity was low.

**Figure 1.**
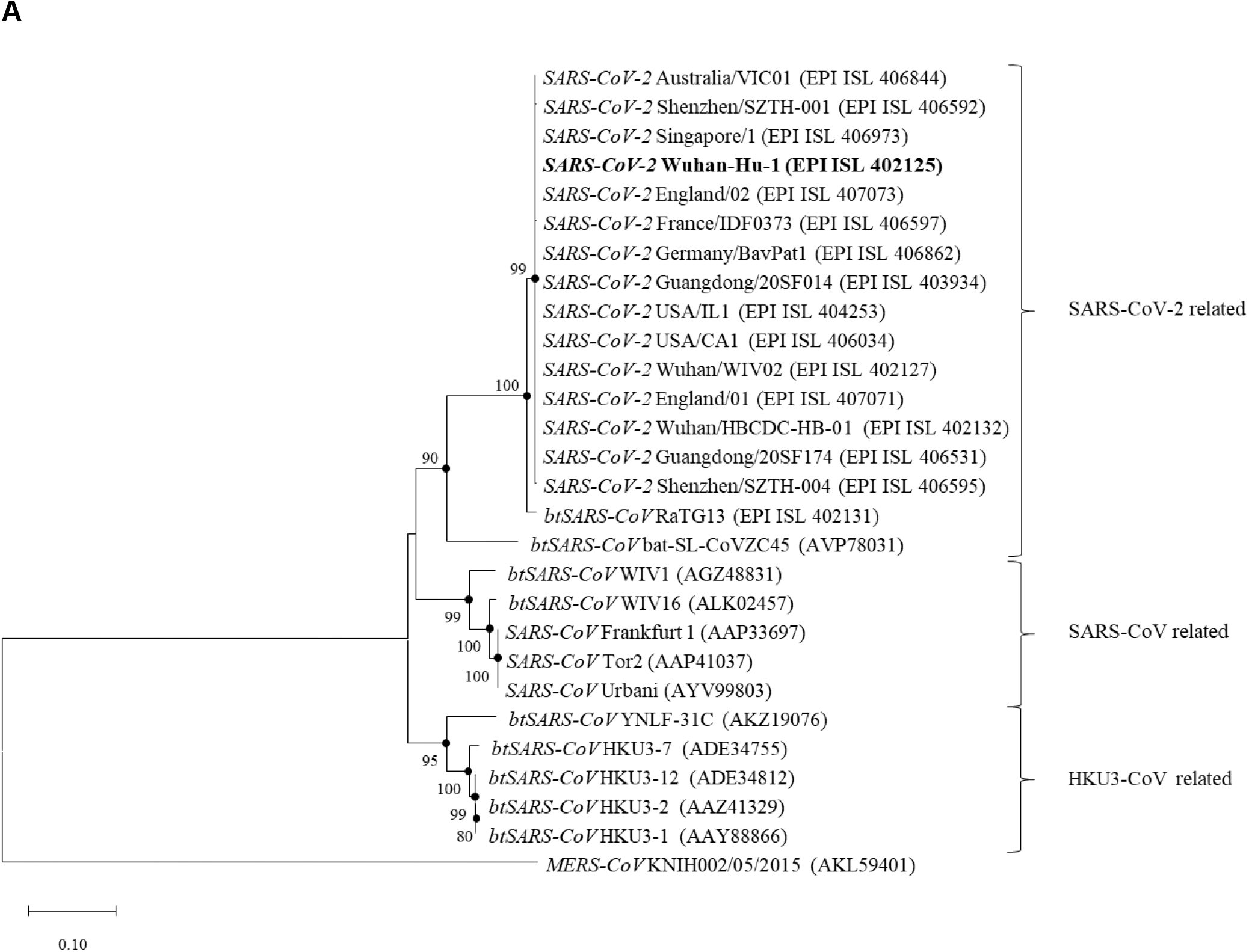
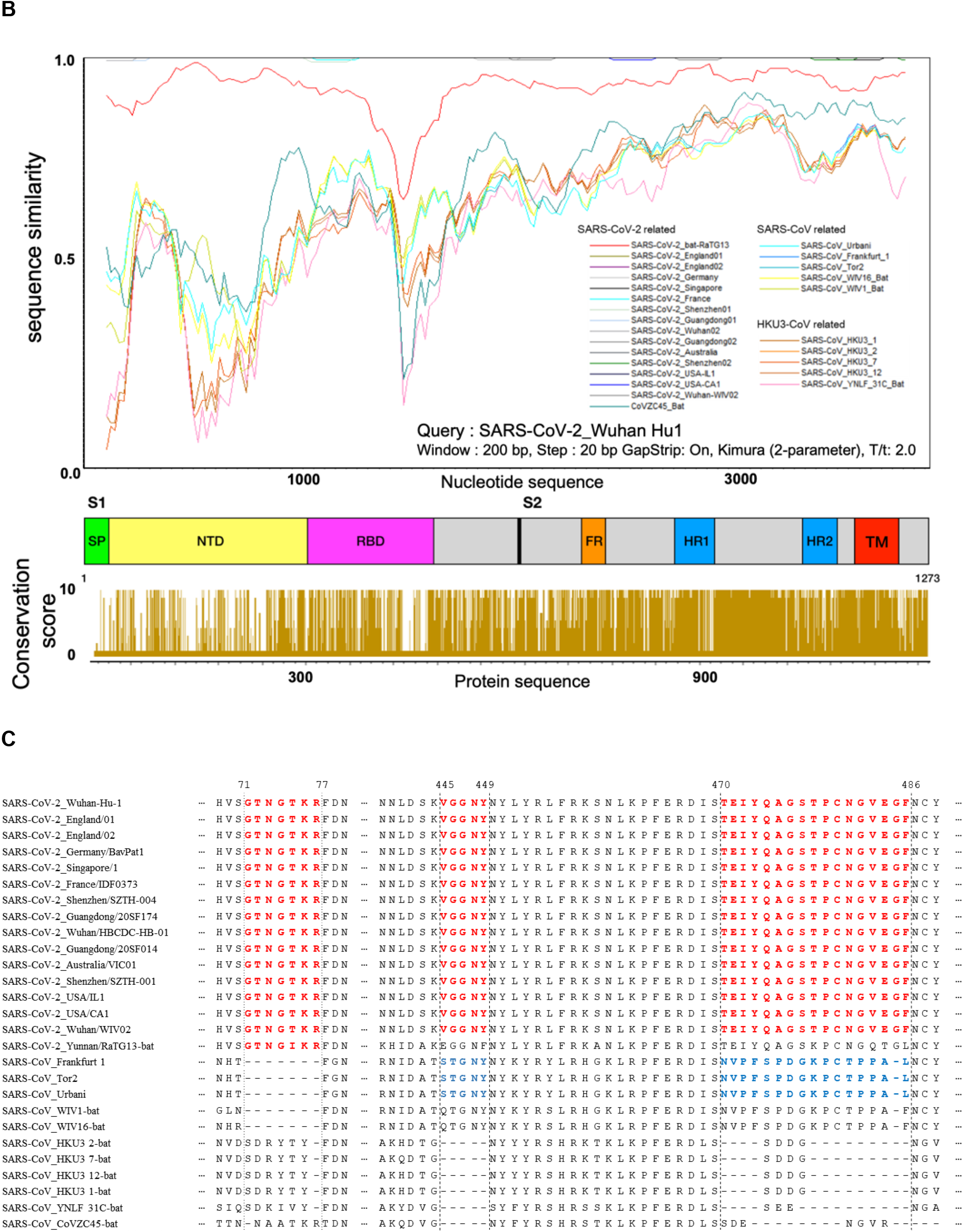
(A) Neighbor-joining tree based on spike protein gene sequences showing the relationship between 2019-nCoV and related SARS-related viruses. The bootstrap value greater than 50 are shown at the branch nodes. The filled circles indicated that the corresponding nodes are conserved in all tree-drawing methods. MERS-CoV KNIH002/05/2015 is used as the outgroup. Bar, 0.1 substitutions per nucleotide position. (B) The upper panel is structure of spike gene of SARS-CoV-2. The middle panel is the result of the Sim plot analysis. The sequence similarity of NTD domain region was the lowest among spike genes. Nevertheless, SARS-CoV-2_bat-RaTG13 had highly conserved in NTD domain than other SARS-CoVs. On the other hand, in the RBD region, not only SARS-CoV but also SARS-CoV-2_bat-RaTG13 showed low sequence similarity. The lower panel indicates the conservation score of the protein sequences of 27 SARS-CoV species. (C) Intersertional regions in SARS-CoV-2 S protein. The amino acid position 445-449 (VGGNY) and 470-486 (TEIYQAGSTPCNGVEGF) were conserved in SARS-CoV-2 related group except bat-SL-CoVZC45 strains (red color), and the corresponding sequences in SARS-related groups were ‘STGNY’ and ‘NVPFSPDGKPCTPPAL’ (blue color).

Interestingly, human pathogenic strains and their closest strains had two insertion sequences in RBD region of SARS-CoV-2. The amino acid position 445-449 (VGGNY) and 470-486 (TEIYQAGSTPCNGVEGF) were conserved in SARS-CoV-2 related group except bat-SL-CoVZC45 strain and the corresponding sequences in SARS-related groups were ‘STGNY’ and ‘NVPFSPDGKPCTPPAL’ (Fig. 1C). The results could not give clear answers that these insertion sequences had directly diverged from the common ancestor of SARS-CoV and SARS-CoV-2 or that the sequences in SARS-CoV-2 were derived from SARS-CoV related group by mobile genetic elements [34]. Nevertheless, the insertion sequences have several antibody epitope regions (Fig. 2) and the two key residues (amino acid position 455 and 486) interacting with human ACE2 [35] which is used as a cellular receptor of btSARS-CoV strain WIV1 [36]. This suggested that these sequence were might be related with human susceptibility and virulence.

**Figure 2.**
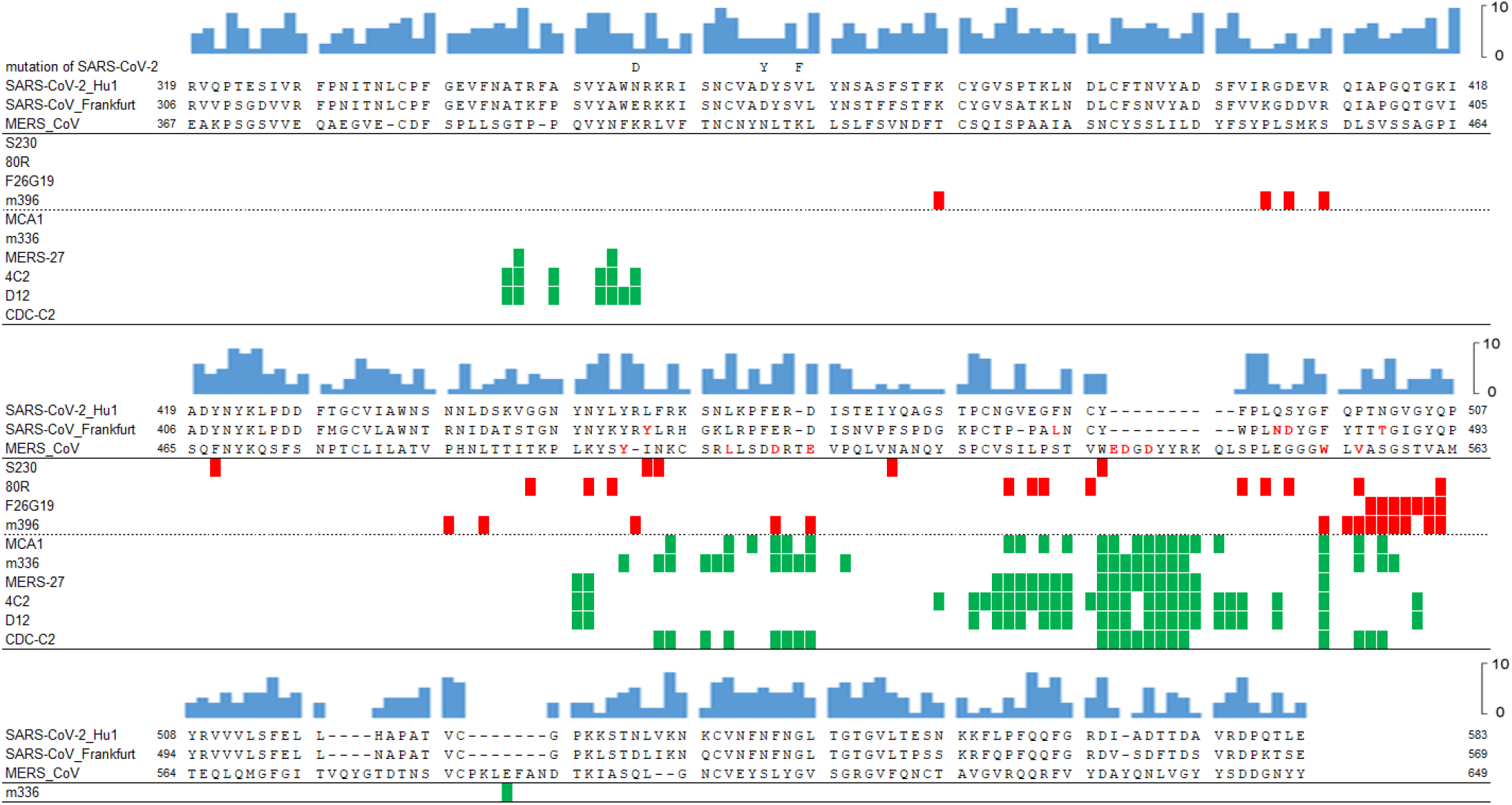
Epitope map and conversation score of the RBD region in SARS-CoV and MERS-CoV S protein. The blue bar chart shows the conservation score. The numbers are the amino acid positions in the S protein. Color boxes indicate binding epitopes for SARS-CoV (red color) and MERS-CoV (green color) antibodies. The amino acid residues in red color indicate binding epitopes for host receptors.

### Identification and analysis of the neutralizing antibody epitopes

Previously, numbers of neutralizing antibodies for SARS-CoV and MERS-CoV have been developed [17, 18]. To suggest possible SARS-CoV-2 neutralizing antibodies, monoclonal antibodies were selected from the literature and PDB (Table 1). Epitope map showed that the antibody-binding residues of S protein are located within RDB region (Fig. 2). Four SARS-CoV neutralizing antibodies had the epitopes about 5 to 14 residues (total 34 residues, average 9.5 residues) of S-RBD and six MERS-CoV neutralizing antibodies bound to 22 to 33 (total 52 residues, average 25 residues) residues. Distribution of the antibody-binding residues indicates that SARS-CoV neutralizing antibodies might be bind to mainly conformational epitopes of S-RDB, whereas MERS-CoV neutralizing antibodies bound to linear epitopes of S-RBD (Fig. 2). Interestingly, the major linear epitope region (EDGDYYRKQL) for MERS-CoV neutralizing antibodies was specific insertion of MERS-CoV S protein. MERS-CoV neutralizing antibodies interacted with three receptor binding residues (E536, D537, D539) in the linear epitope region, which results in the neutralizing activity of antibodies by directly interferes the binding between S protein and dipeptidyl peptidase 4 of human . In addition, the difference of binding aspect of neutralizing antibodies might be caused by the difference of subdomain structure of receptor binding motif (RBM). The RBM of SARS-CoV S protein is made of mainly coiled structure with two short β-sheets, whereas the RBM of MERS-CoV S protein consists of four long β-sheets [37]. Sequence alignment revealed that RBD of SARS-CoV-2 was more similar to that of SARS-CoV than MERS-CoV (Fig. 2). Therefore, this suggested that SARS-CoV neutralizing antibodies could be effective for SARS-CoV-2.

### S-RBD structure modeling and superimposition of neutralizing antibodies

Human infection of SARS-CoV-2 was firstly reported in Wuhan, Hubei province, China last December [4]. Previous studies have reported several results for the interaction between S protein of SARS-CoV-2 and ACE2 as a receptor [13, 14, 16, 38]. However, any interaction of SARS-CoV-2 S protein with developed neutralizing antibodies for SARS-CoV and MERS-CoV has not been reported yet. The structure of SARS-CoV-2 S protein was used S protein which revealed by cryo electron microscopy structure. Subsequently, the missing region of SARS-CoV-2 S-RBD region comprising of 181 amino acids were built from SARS-CoV S proteins (PDB ID: 6NB7) which were good structural templates (Fig. 3 box). In the S-RBD structure, we also displayed experimentally defined epitope information based on position specific aligment with SARS-CoV or MERS-CoV antibody binding epitopes.

**Figure 3.**
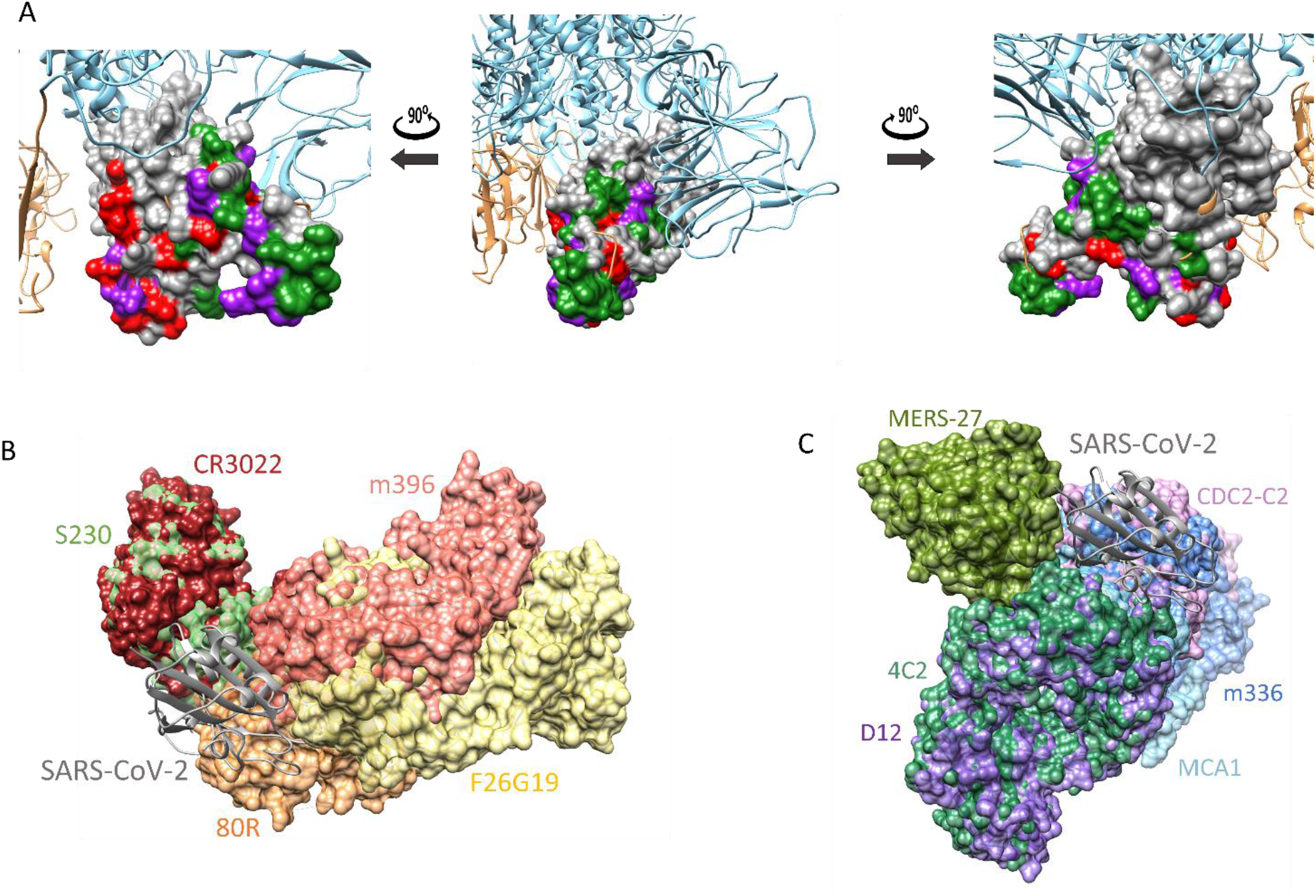
The three dimensional structure of SARS-CoV-2 S protein. (A) The three dimensional structure of RBD domain in S protein was colored gray. The RBD structure shown from various sides. On the surface representation, the SARS-CoV antibodies, MERS-CoV antibodies, and both SARS-CoV and MERS-CoV antibodies binding epitopes are colored red, green, and purple, respectively. The sky blue color represents SARS-CoV-2 S protein and the RBD domain were highlighted with orange color. The red box indicates the RBD region, which is containing SARS-CoV or MERS-CoV antibody binding epitope. The predicted RBD structure of SARS-CoV-2 S protein in complex with five SARS-CoV antibodies (B) and six MERS-CoV antibodies (C). The complex structure was predicted by integrating the previously known complex structures of SARS-CoV or MERS-CoV with antibodies using the superimposition of structures. Each colored structure in surface representation indicates antibody labeled with the same color. More detail information about antibodies were described in Table 1.

To visualize the overall antibody binding region to SARS-CoV-2, we superimposed the predicted structure of SARS-CoV-2 RBD protein at the X-ray crystal structure of known antibody-antigen complex from SARS and MERS (Fig. 3). The structures of five antibodies including m396, 80R, F26G19, S230, and CR3022 developing to prevent SARS-CoV were aligned on SARS-CoV-2 S-RBD successfully (Fig. 3A). The six MERS-CoV antibodies such as MERS-27, CDC2-C2, m336, 4C2, D12, and MCA1 were also aligned on SARS-CoV-2 S-RBD (Fig. 3B). Because the X-ray crystal structure of CR3022 was not revealed, the optimized structure was predicted using antibody homology modeling by 1000 structures generated using Rosetta program. As a results, two SARS-CoV antibodies including CR3022 (−13.91 dG score) and F26G19 (−15.98 dG score) and MERS-CoV D12 (−14.01 dG score) antibody had higher binding score than other antibodies with SARS-CoV-2 S-RBD region. However, various MERS-CoV antibodies did not match SARS-CoV-2 because MERS-CoV antibodies were interacted with the outter regon of S-RBD which was located in major linear epitope region (EDGDYYRKQL) (Fig. 2).

### Comparison of Antibody-RBD protein binding interaction

Based on antibody-antigen docking simulation, we calculated the binding scores between 11 antibodies and S-RBD structures. The antibody-antigen docking simulation generated not only the crystal structures of SARS-CoV and MERS-CoV S-RBD proteins, but also the high-quaility homology models with SARS-CoV-2 S-RBD. To suggest S-RBD binding antibody, antibody-RBD docking comparisons were performed using the mean value of caculated scores from the generated models. The mean scores of the docking simulation are shown in Table 2.

**Table 2.** Antibody-antigen docking score and experimental affinity

| Antibody | SARS/MERS-CoV S-RBD |  | SARS-CoV-2 S-RBD |  | p-value |
| --- | --- | --- | --- | --- | --- |
|  | Docking (dG score) | Experiment (nM) | Docking (dG score) | Experiment (nM) |  |
| m396 (SARS) | -21.92 | 0.0046 | -6.92 | No binding | < 2.2e-16 |
| 80R (SARS) | -12.04 | 1.59 | -3.59 | - | 3.986e-07 |
| S230 (SARS) | -7.18 | - | -7.48 | - | 0.7731 |
| F26G19 (SARS) | -17.16 | 0.45 | -15.98 | - | 2.049e-05 |
| CR3022 (SARS) | -11.21 | 0.125 | -13.91 | 6.28 | 0.00367 |
| MERS27 (MERS) | -13.12 | 71.2 | -4.26 | - | < 2.2e-16 |
| CDC2-C2 (MERS) | -28.21 | 2.65 | -11.63 | - | < 2.2e-16 |
| m336 (MERS) | -23.21 | 0.0994 | -12.05 | No binding | < 2.2e-16 |
| MCA1 (MERS) | -26.74 | - | -10.69 | - | < 2.2e-16 |
| 4C2 (MERS) | -21.12 | 317 | -11.48 | - | < 2.2e-16 |
| D12 (MERS) | -22.03 | 6.63 | -14.01 | - | < 2.2e-16 |

Among the SARS-CoV antibodies, only CR3022 showed that the binding affinity of SARS-CoV-2 was higher than SARS-CoV. In addition, the docking score distribution of CR3022 was significantly changed between SARS-CoV-2 and SARS-CoV-2 (Fig. 4). For the CR3022 antibody, the mean score of binding affinity was increased from −11.21 dG score (SARS-CoV, crystal structure) to −13.91 dG score (SARS-CoV-2, cryo-EM structure) with a *p-value* of 0.00367. The binding affinity of all antibody-antibgen docking was tested using 1000 generated structures. Interestingly, the CR3022 was experimentally performed for the binding effect of SARS-CoV-2 S-RBD [14]. The researchers found that the CR3022 had the binding effect anainst SARS-CoV-2 S-RBD and that m396 and m336 antibodies did not bind to SARS-CoV-2 S-RBD. The researchers also reported that the binding affinity of CR3022 was increased to 6.28nM with SARS-CoV-2 from 0.125nM with SARS-CoV, The results of the docking simulation were consistent with the evidence although more research was needed to prevent effects, including an experiment using live viruses.

**Figure 4.**
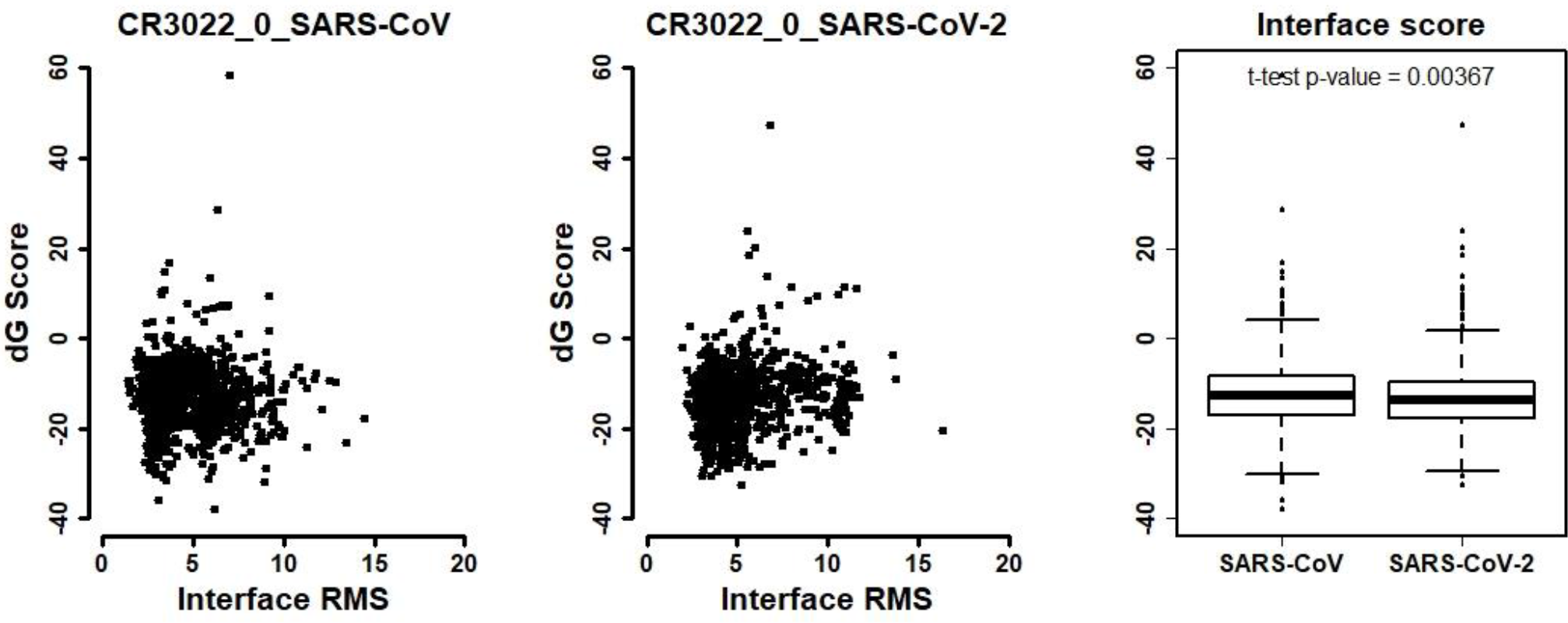
The distiribution of docking scores between antibody and antigen using interface RMD (interface RMS) versus the binding energy (dG binding). The docking simulation were performed by 1000 generated antibody-antigen structures. The statistical significance was tested using student’s *t-test*. (A) Mean score of docking simulation between CR3022 and SARS-CoV S-RBD was −11.21 dG score, and mean score of CR-3022 and SARS-CoV2 S-RBD was −13.61 dG score (*p-value* = 0.00367).

## Conclusions

The fact that CoVs similar to SARS in Chinese bats is most identical to SARS-CoV-2 suggests that SARS-CoV may have been originated from a common ancestral bat CoV. Comparing the sequences among the three groups, various deletions were observed in the SARS-CoV related group. Especially, amino acid positions 71-77 (GTNGTKR) in the NTD region of the S protein, 445-449 (VGGNY) and 470-486 (TEIYQAGSTPCNGVEGF) were noteworthy. The regions were highly conserved in SARS-CoV-2, unlike other SARS-CoVs.

Among the neutralizing antibodies for SARS-CoV and MERS-CoV, CR3022 was predicted to have better binding affinity to the S-RBD region of SARS-CoV-2 than other antibodies. The comparison of antibody binding region between SARS-CoV-2 and other coronaviruses, such as SARS-CoV and MERS-CoV, was conducted to apply the suitable diagnostic or therapeutic antibodies and vaccines that are mimetics of extremely infectious SARS-CoV-2.

## Acknowledgment

We thanks to Prof. Jason S. McLellan for providing cyro-EM structure of of SARS-CoV-2 S protein, Korea Institute of Science and Technology Information’s 5th supercomputer NURION, and bio bigdata center of Clinomics Inc.. This work was supported by National Research Council of Science and Technology grant by the Ministry of Science and ICT (Grant No. CRC-16-01-KRICT).

